# Mustache: Multi-scale Detection of Chromatin Loops from Hi-C and Micro-C Maps using Scale-Space Representation

**DOI:** 10.1101/2020.02.24.963579

**Authors:** Abbas Roayaei Ardakany, Halil Tuvan Gezer, Stefano Lonardi, Ferhat Ay

## Abstract

We present Mustache, a new method for multi-scale detection of chromatin loops from Hi-C and Micro-C contact maps. Mustache employs scale-space theory, a technical advance in computer vision, to detect blob-shaped objects in a multi-scale representation of chromatin contact maps parametrized by the size of the smoothing kernel. When applied to high-resolution Hi-C and Micro-C data, Mustache detects loops at a wide range of genomic distances, identifying potential structural and regulatory interactions that are supported by independent conformation capture experiments as well as by known correlates of loop formation such as CTCF binding, enhancers and promoters. Unlike the commonly used HiCCUPS tool, Mustache runs on general-purpose CPUs and it is very time efficient with a runtime of only a few minutes per chromosome for 5kb-resolution human genome contact maps. Extensive experimental results show that Mustache reports two to three times the number of HiCCUPS loops, which are reproducible across replicates. It also recovers a larger proportion of published ChIA-PET and HiChIP loops than HiCCUPS. A comparative analysis of Mustache’s experimental results on Hi-C and Micro-C data confirms strong agreement between the two datasets with Micro-C providing better power for loop detection. Overall, our experimental results show that Mustache enables a more efficient and comprehensive analysis of the chromatin looping from high-resolution Hi-C and Micro-C datasets. Mustache is freely available at https://github.com/ay-lab/mustache.

## Background

Recent studies have revealed that chromatin has a well-organized structure in the eukaryotic nucleus, which is highly regulated in accordance to the stage of the cell cycle, environmental cues and disease conditions [1, 2, 3, 4]. In turn, the 3D structure of the chromatin plays a critical role in many essential cellular processes, including the regulation of gene expression and DNA replication [5, 6, 7]. Therefore, it is of great importance to systematically study the chromatin organization to understand how folding properties and looping events influence the cell-specific biological functions of distinct regulatory regions of the DNA. To date, Hi-C has been the main assay of choice for discovering genome-wide chromatin contacts/interactions [8, 9]. More recently, Micro-C [10], which replaces the restriction enzyme in Hi-C with micrococcal nuclease for digestion, enabled the generation nucleosome resolution chromosome folding maps of mouse and human cells [11, 12]. With the decreasing cost of sequencing and optimization for smaller cell numbers, both Hi-C and Micro-C assays are expected to produce much higher resolution reference contact maps for a diverse set of organisms and cell lines [9, 13, 12].

Chromatin loops are defined as pairs of genomic sites that lie far apart along the linear genome but are brought into spatial proximity by a mechanism called *loop extrusion* [14, 15, 16]. Several methods have been developed to detect chromatin loops or statistically significant/enriched chromatin interactions from Hi-C and Micro-C contact maps [17, 9, 18, 19]. These existing methods broadly fall into two groups. The first group, which we call *global enrichment-based methods*, contains methods that *(i)* globally fit statistical/probabilistic models to the contact map data and *(ii)* assign *p*-values to each individual pixel/entry in the contact map by comparing the observed count values to the expected values computed from the fitted model. For instance, Fit-Hi-C [17] uses a monotonic spline to model the relation between contact probability and the genomic distance of the interacting loci, then estimates the statistical confidence of each contact with respect to this expectation and a coverage-based correction factor, using a binomial distribution. In a similar method, HiC-DC [18] estimates the statistical significance of chromatin contacts from Hi-C experiments using a hurdle negative binomial regression to account for both the zero inflation and over-dispersion of contact counts as well as systematic sources of variation in Hi-C read counts such as distance-dependent random polymer ligation, GC content and mappability bias. Several crucial drawbacks common to these two methods, and to any other method that uses only a global background [20], are: *(i)* the locality information in the contact map is not taken into account in the modeling, *(ii)* each pixel/contact count is considered independent of its surrounding pixels, *(iii)* pixels that are in the vicinity of a strong loop are also deemed statistically significant with respect to the global background (i.e., bystander effect). As a consequence, a large number of reported significant contacts are likely to cluster around a few very strong pixels or loops, making it difficult to interpret the large number of direct and indirect enrichment lumped together. Recent variations of these methods employ post filtering strategies for discarding calls that are likely to be indirect interactions [21, 22], however, for deeply sequenced Hi-C data, the number of resulting calls still remains very large (e.g., in the order of hundreds of thousands). Therefore, the global enrichment-based methods, even though are important for discovering preferential enrichment of proximity among functional regulatory elements such as promoters and enhancers [22], are not ideal if only the strongest and structural loops, such as those demarcating domain boundaries and in between convergent CTCF binding sites, are of interest [9].

The second group of methods, which we name *loop callers*, identify 2D peaks in contact map that are not only significantly higher than expected from the global background, but also are *local maxima* with respect to their neighboring pixels. For example, Rao *et al.* developed a method called Hi-C Computational Unbiased Peak Search (HiCCUPS) that detects chromatin loops in deeply sequenced high-resolution Hi-C maps [9]. HiCCUPS examines each pixel (locus pair) in the contact map by comparing its contact frequency to that of neighboring pixels. More specifically, HiCCUPS identifies loops by finding “enriched” pixels, that is, locus pairs whose contact counts are significantly higher than that of (1) pixels to its lower-left, (2) pixels to its left and right, (3) pixels above and below, and (4) a doughnut-shaped region surrounding the pixel of interest. This approach typically produces a small set of loop calls (∼10k) where each loop corresponds to a clear enrichment in the contact map. However, HiCCUPS and other methods in this class use a fixed representation of the contact map and a fixed-size local neighborhood to model the background intensities. Therefore, chromatin loops that are caused by proximity of larger (or smaller) regions of DNA and subsequently reflected as larger (or smaller) blobs in the contact map, as compared to the scale of the fixed representation, may not meet the local filtering criteria and will not be reported.

In this paper, we present a new loop caller named Mustache that addresses the drawbacks mentioned above. Mustache uses the scale-space representation of a contact map to model and identify chromatin loops at multiple resolutions. Mustache utilizes a set of carefully designed filters to report only locally-enriched pixels as loops. Our experimental results show that Mustache detects chromatin loops that are reproducible, have high support from aggregate peak analysis and are independently supported by other conformation capture experiments as well as by genomic and epigenomic correlates of loop formation. Given the orders of magnitude of difference in the resulting calls from Mustache and global enrichment-based methods such as Fit-Hi-C and HiC-DC (e.g., 20k vs 700k vs 1M, respectively), here we focus on comparing Mustache to the most commonly used loop calling method with local filtering, which is HiCCUPS. Our comparisons show that Mustache provides better statistical power in detecting loops compared to HiCCUPS while detecting the majority of HiCCUPS loops. We present several lines of evidence suggesting that the additional loops detected only by Mustache are not false-positives, but are rather *bona fide* looping events with visible enrichments in contact maps. We also show Mustache’s efficacy for Micro-C data and consistency of its loop calls from Micro-C and Hi-C contact maps of the same cell line [12]. Based on the results presented here, we believe that Mustache will become an essential tool in the analysis of high-resolution Hi-C and Micro-C contact maps, which are being produced in large numbers by the 4D Nucleome project and other efforts [23].

## Methods

### Scale-Space modeling

Objects in real world, as opposed to mathematically-defined abstract entities such as points and lines, are composed of a variety of structures and textures at different scales which often makes them difficult to detect in the absence of *a priori* knowledge about their true scales. A way of addressing this challenge is to describe each object at multiple scales. In the specific problem on contact maps we are interested here, significant chromatin interactions are “blob-shaped objects” with a scale that depends on their size and other properties of the interacting genomic regions (e.g., CTCF binding, presence of regulatory elements).

Scale-space theory is a framework developed by the Computer Vision community for multi-scale representation of image data. In scale-space theory, each image is represented as a set of smoothed images. In order to build a scale-space representation of an image, a gradual smoothing process is conducted via a kernel of increasing width, producing a one-parameter (i.e., kernel size) family of images. This multi-scale representation makes it possible to detect smaller patterns at finer scales, while allowing the detection of larger patterns at coarser scales (Figure 1a). The most common type of scale-space representation uses the Gaussian kernel because of its desirable mathematical properties. In particular, the causality property of Gaussian kernel guarantees that any feature at a coarse resolution scale is caused by existing features at finer resolution scales. This property makes sure that the smoothing process cannot introduce new extrema in the coarser scales of the scale-space representation of an image [24], which is critical for the problem we tackle here.

**Figure 1:**
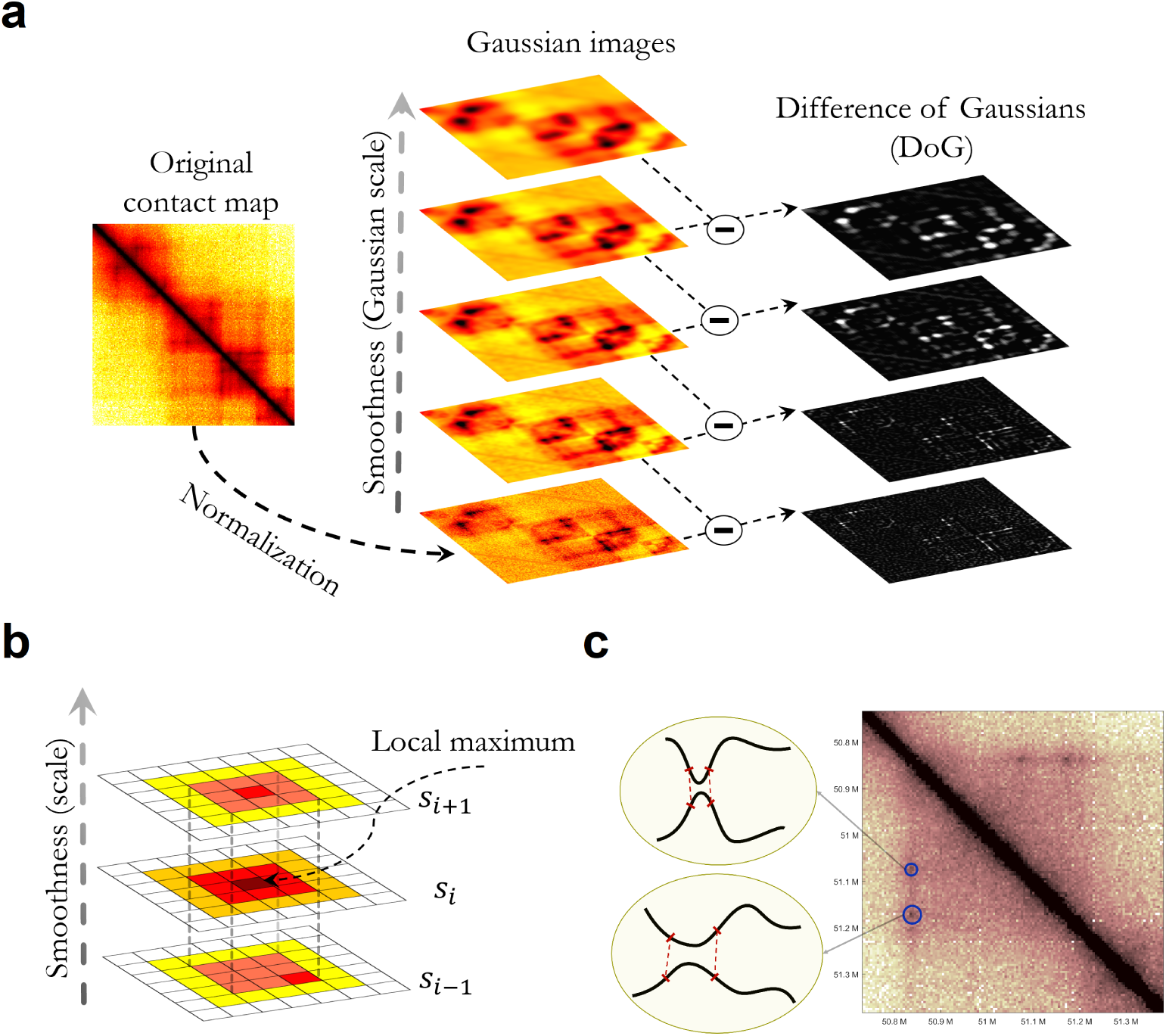
**(a)** The initial contact map is repeatedly convolved with increasing 2D Gaussians to produce a scale-space representation of the image (shown on the left). Pairwise adjacent Gaussian images are subtracted to produce the difference-of-Gaussian (DoG) images (on the right); **(b)** Maxima of the difference-of-Gaussian images are detected by comparing each pixel to its 3×3×3 neighborhood in (*x, y, σ*) space. Note that DoG is a local maximum at (*x, y*) location at scales *s*_*i*_ and *s*_*i*+1_ but not at scale *s*_*i*−1_, therefore passing the first filtering step criteria; **(c)** Chromatin loops can be caused by the contact between pairs of DNA segments at different scales.

The Gaussian-kernel scale-space representation of an image *A*(*x, y*) is a function *L*(*x, y, σ*) obtained from the convolution of a variable-scale Gaussian *G*(*x, y, σ*) with the input image, as follows

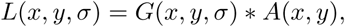

where * represents the convolution operation in *x* and *y*, and

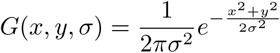

is a 2D Gaussian (see [25] for more details).

Blob-shaped objects can be typically detected in an image by finding the strong responses in the application of the *Laplacian of the Gaussian* operator with an image, as follows

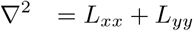

Lindeberg showed that the normalization of the Laplacian with the factor *σ*^2^∇^2^ provides the scale invariance required for detecting blob-shaped objects at different scales [24]. According to Lowe [26], the scale-normalized Laplacian *σ*^2^∇^2^ can be accurately and efficiently estimated by the *difference-of-Gaussian* (DoG) function. Therefore, blob-shaped objects of varying scale can be detected from the scale-space maxima of the DoG function *D*(*x, y, σ*) convolved with the image, which can be computed from the difference of two nearby scales (in a scale-space representation) separated by a constant multiplicative factor *k*, as follow

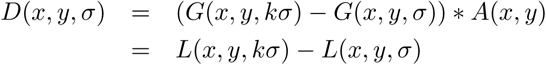

### Development of Mustache

Since a chromatin contact map can be represented by a digital image, we assume that a chromatin loop can be described as a somewhat-circular (blob-shaped) object with its own specific scale (that can be determined using the scale-space representation). Thus Mustache’s objective is to find blob-shaped regions of interactions with high statistical significance, i.e., regions with an average interaction significantly greater than expected. Due to random polymer interactions driven by one-dimensional genome proximity, interactions between pairs of loci that are closer in genomic distance are more frequent than interactions between loci at higher genomic distances. To account for the amplification of contact frequency due to 1D proximity, Mustache performs a simple local *z*-normalization of the interaction frequencies in the contact map *A* with respect to their genomic distances along each diagonal *d*. Then, Mustache re-scales the interactions by the logarithm of the expected interaction of the corresponding distance, as follows

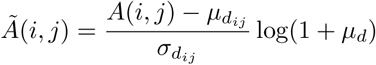

where *d* = |*j* − *i*|, *µ*_*d*_ is the average interaction on diagonal *d*, and 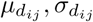 (not to be confused with Gaussian scale *σ*) are the local average and standard deviation along the diagonal *d* in a ±1Mb neighborhood, respectively. Then, Mustache constructs the scale-space representation *D* of the normalized contact map *Ã*. As explained above, in order to compute *D*(*x, y, σ*), Mustache convolves *Ã* with Gaussians that have increasing scales (i.e., *σ, kσ, k*^2^*σ*, …). This process produces a set of smoothed contact maps separated by a constant factor *k* in scale-space. Mustache computes the difference of Gaussians (DoG) by subtracting pairwise adjacent smoothed contact maps (Figure 1a).

Mustache uses two octaves of scale-space, each divided into *s* intervals, such that *k* = 2^1*/s*^ [25]. Mustache computes the *p*-value 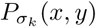 for each pixel *D*(*x, y, σ* = *σ*_*k*_) by fitting a Laplace distribution on each scale of DoG, as follows

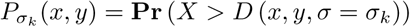

where *X* is distributed according to the Laplace distribution. After computing the difference of Gaussians *D*(*x, y, σ*), Mustache searches for local maxima in the 3D space over the parameters *x, y* and *σ*. Specifically, Mustache defines a 3D local maximum as a pixel (*x, y, s*_*i*_) (where *s*_*i*_ is a specific scale) that is greater than all its neighboring pixels at scale *s* (eight surrounding pixels) as well as its nine neighboring pixels in the scale above (*s*_*i*+1_) and nine neighboring pixels in the scale below (*s*_*i*−1_) within a 3 × 3 × 3 cube, as illustrated in Figure 1b. Such local maxima are selected as candidate loops at scale *s* and the rest are discarded (non-maxima suppression). In case a pixel is a local maximum at multiple non-adjacent scales, Mustache reports the minimum *p*-value across all scales for that specific pixel as its significance.

In order to find high-confidence and locally-enriched loops, the set of detected candidates undergoes a few additional filtering steps. In the first step, Mustache removes candidates that are not local maximum in at least two consecutive scales in a 3 × 3 two-dimensional neighborhood (i.e., it discards candidates that are local maximum at scale *s*_*i*_ but not in *s*_*i*−1_ and not in *s*_*i*+1_). Figure 1b illustrates an example candidate pixel that passes the non-maxima suppression step (being a maximum in a 3×3×3 neighborhood) as well as the first filtering step. Observe that the pixel at (*x, y*) location (center pixel) is a local maximum in its 3 × 3 neighborhood at scales *s*_*i*_ and *s*_*i*+1_ but not a local maximum at scale *s*_*i*−1_. In the second filtering step, Mustache finds connected components using 8-connectivity (i.e., a 3 × 3 neighborhood around each pixel) on a binary matrix in which an entry is set to one when that pixel is a candidate loop at any scale. For each connected component, Mustache reports the single representative pixel that has the lowest *p*-value. In the third filtering step, Mustache filters out candidates that are located in sparse regions of the contact map. Specifically, it discards pixels whose neighborhood, as defined by a size equal to the *σ* of the scale the candidate was detected from, contains more than 20% pixels with zero count in the raw Hi-C contact map. Such pixels are likely enriched near repetitive and unmappable regions of the genome and may introduce false positives. In the fourth and final filtering step, candidates with contact count smaller than two times the expected count for that specific diagonal (i.e. the average contact count of all pairs with that corresponding genomic distance) are discarded. The remaining set of pixels are reported as loop calls together with their statistical significance and the scale of the Gaussian that the reported *p*-value was identified from. Mustache uses the Benjamini-Hochberg [27] procedure to correct values for multiple hypothesis testing.

## Results

### Mustache detects chromatin loops from publicly available contact maps

We ran Mustache on Hi-C contact maps for human cell lines GM12878 and K562 obtained from Rao *et al.* [9], and Hi-C and Micro-C contact maps of HFFc6 cell line obtained from [12], all at 5kb resolution (Supplementary Table 1). For assessing the consistency of loop calls at different resolutions and for measuring reproducibility across replicates, we also used 10kb GM12878 Hi-C contact maps. All contact maps were produced using a minimum read alignment quality MAPQ≤30. GM12878, K562 and HFFc6 Hi-C data consisted of approximately 4.9B, 1B and 3B read pairs, respectively. HFFc6 Micro-C data had over 4B reads. For all experiments, we considered contacts within the genomic distance range of 20kb to 2Mb. We selected these datasets because they are the most extensively studied, they are generated with very high-depth sequencing and these cell lines have readily available orthogonal datasets including ChIP-seq, ChIA-PET and HiChIP, which we used for validating our results. For the 5kb resolution Hi-C map of GM12878 (obtained by combining the two replicates), we employed Mustache using two octaves with default parameters (*σ*_0_ = 1.6 and *s* = 10) resulting in 18, 068 loop calls for q-value < 0.05. For the 10kb resolution GM12878 data we obtained 14,045 loops from the combined contact map, and 11, 872 and 10, 976 loops from the primary and replicate experiments, respectively, using the same q-value threshold of 0.05. For the combined 5kb K562 Hi-C data from the same publication, we identified 8, 975 loops at a q-value of 0.1. On the HFFc6 cell line, Mustache called 16, 132 loops from the Hi-C data and 36, 494 loops from Micro-C data with the q-value < 0.01, both at 5kb resolution. Using a more stringent value of < 0.001 on the Micro-C data, Mustache detected 24, 045 loops. We selected the false discovery rate (FDR) thresholds such that the number of loop calls were comparable across different assays (i.e., Hi-C versus Micro-C) or were comparable across different cell types. In order to carry out a fair comparison with HiCCUPS, we selected the top-*k* Mustache loops with respect to their statistical significance, where *k* was set to match the number of loops called by HiCCUPS on the same contact map.

### Comparison of loop calls from Mustache and HiCCUPS on Hi-C data

As discussed, methods that detect enrichment of contact counts with respect to a global background model tend to report a number of significant contacts a few orders of magnitude higher (over 700k and 1M for GM12878 Hi-C data) than loop callers with local filters such as HiCCUPS (∼10k) and Mustache (∼20k) on the same data. Even though the global background models are valuable for finding potential interactions among functional elements such as enhancers and promoters, most of the detected enrichments do not correspond loop anchors demarcating domain boundaries and/or specific loops between convergent CTCF binding sites which are argued to occur due to loop extrusion [16, 14, 9, 15]. Therefore, we focused our comparative analysis with what we call here as “loop calling” methods, which are geared towards detecting the strongest and domain-demarcating pixels, such as the commonly used HiCCUPS method [9]. Using the publicly available Hi-C contact maps described above, we conducted several comparisons between Mustache and HiCCUPS loop calls, which were obtained directly from the GEO entry for that work [9]. First, we performed a genome-wide comparison of Mustache results against 9, 448 GM12878 and 6, 057 K562 loop calls from HiCCUPS. To compute the overlap between the two methods, we defined two loop calls as *matched* if the ±5kb area around the center of each loop overlapped each other (inclusive of corners and edges). For this analysis all loop calls by Mustache were reported at 5kb resolution, whereas HiCCUPS reported a mix of loops at 5kb and 10kb. Figure 2a-b illustrates that there was a good agreement between the two methods. Mustache recovered nearly 81% HiCCUPS loops in GM12878 (Figure 2a) and 65% of HiCCUPS loops in K562 (Figure 2b). When we compared loop calls from GM12878 to that of K562, we observed that Mustache and HiCCUPS reported a similar fraction of common loops suggesting that the two methods have similar cell type specificity **(Supplementary Figure 1)**.

**Figure 2:**
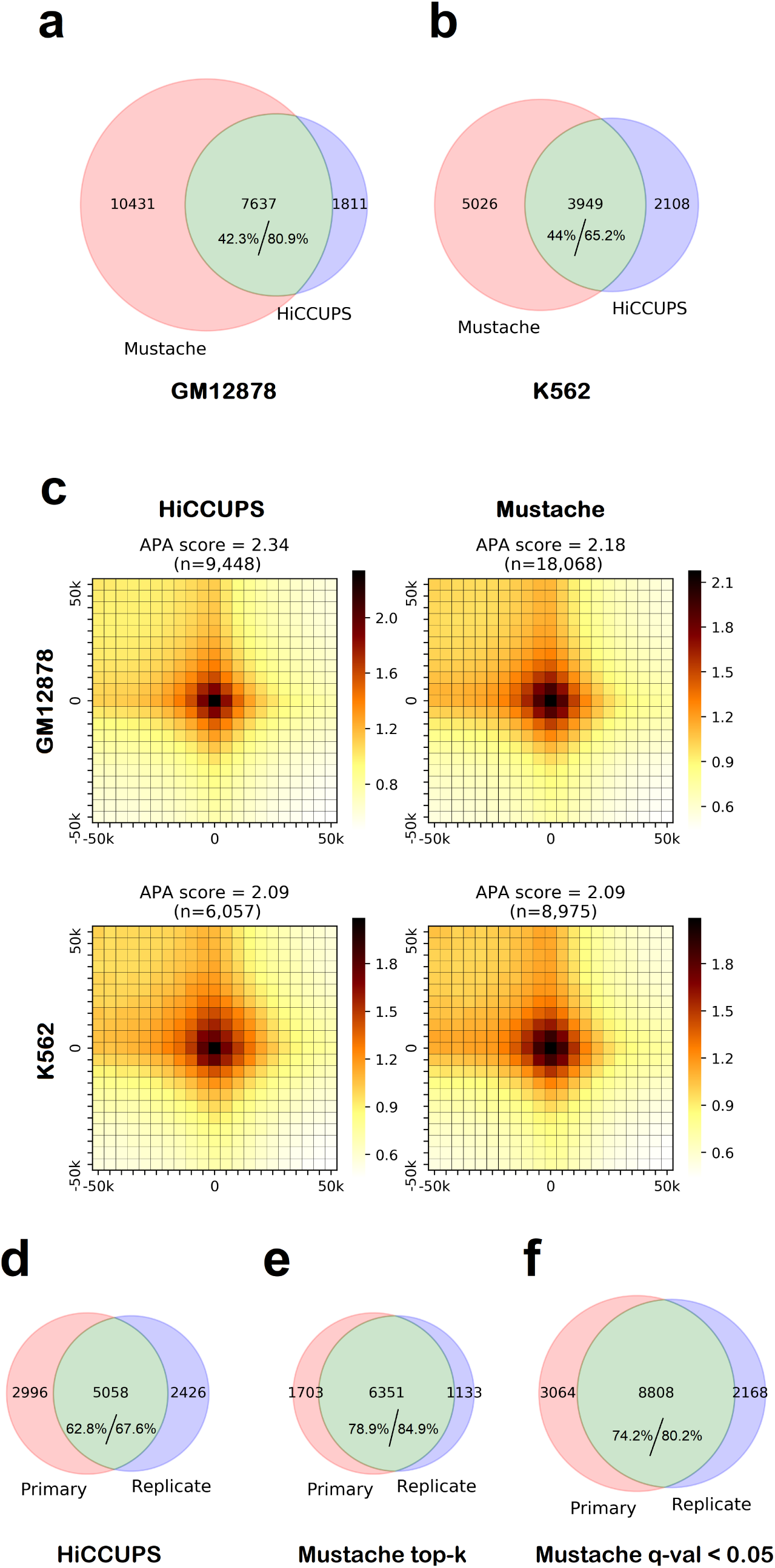
Comparison of chromatin loops detected by Mustache and HiCCUPS in GM12878 and K562 cell lines. The agreement between HiCCUPS and Mustache loops are shown as Venn diagrams for replicate-combined Hi-C contact maps of **(a)** GM12878, and **(b)** K562 cell lines. The overlap between the two loop sets is shown in green and the percentages of overlap with respect to each set are reported separately; **(c)** APA plots for Mustache and HiCCUPS loops in GM12878 and K562 cell lines. The APA score for the enrichment of center is reported above each plot; **(d)** The overlap between HiCCUPS reported loops on two replicates of GM12878; **(e)** The overlap between Mustache reported loops on two replicates of GM12878 when we restrict the number of loops to match that of HiCCUPS for each replicate (top-*k* setting); **(f)** The overlap between Mustache reported loops on the two replicates for a *q*-value threshold of 0.05. For **(d-f)**, when replicates are analyzed separately, we use 10kb resolution Hi-C contact maps.

Figure 2a-b shows that Mustache reported a substantial number of additional loops for each cell type compared to HiCCUPS. In order to further evaluate this difference, we quantified how well Mustache loops were supported by the Hi-C data compared to HiCCUPS using aggregate peak analysis (APA) [9, 28]. To generate APA plots, we aggregated contact counts over all detected HiCCUPS loops and top-*k* Mustache loops (equal to that reported by HiCCUPS) between 150kb to 1Mb and their ±50kb neighborhood. The result is illustrated as a 21 × 21 heatmap (at 5kb resolution) in which darker color indicates higher contact count. A strong dark pixel at the center of the heatmap indicates a specific enrichment of Hi-C contacts with respect to local background for the detected loops. The enrichment is also quantified by the APA score, which is the ratio of the value of the center pixel to the average value of pixels 15 − 30kb upstream and downstream [28]. Figure 2c shows APA plots and APA scores on GM12878 and K562 cell lines for all reported HiCCUPS loops and a matching number of the top-*k* Mustache loops. These results show that loops from both methods exhibited comparable enrichment with a slightly more localized enrichment for HiCCUPS in GM12878 and for Mustache in K562 contact maps. For assessing differences in the cell type-specificity of these loop calls, we conducted a reverse APA analysis where loops were called from one cell type and the aggregated counts were gathered from the other. This analysis showed enrichment of center pixels, albeit lower than the matched cell type case, for both methods but no significant differences between them in the amount of that enrichment **(Supplementary Figure 2)**.

To measure the agreement between Mustache and HiCCUPS loop calls from replicate experiments, we compared them on two replicates of GM12878 cell line. For Mustache, we either used a fixed *q*-value threshold (0.05) or the top-*k* loops where *k* was set to match the number of reported HiCCUPS loops. Since HiCCUPS calls for individual replicates were at 10kb resolution, Mustache was also run on 10kb resolution replicate-specific maps. For assessing whether two loops match each other, one from each replicate, we used the overlap of ±5kb area around the center of each loop as previously described. Figure 2d-f show that while both methods have a considerable overlap between replicates, Mustache reported over a thousand more (reproducible) loops for the same number of calls and over 75% more (5, 058 vs 8, 808) reproducible calls for the fixed significance threshold of q-value < 0.05. These results indicate that Mustache did not compromise the reproducibility in either scenario and had better self-consistency compared to HiCCUPS. The comparative analysis of the distribution of genomic distances for reported loops confirmed that the higher reproducibility of Mustache results cannot be attributed to preference in reporting shorter or longer distance loops as the the two methods lead to virtually identical distance distributions (**Supplementary Figure 3**).

### Mustache identifies additional loops with enrichment in expected features of chromatin looping

In the following experiment, we evaluated the convergence of CTCF sites on the anchors of chromatin loops detected by Mustache and HiCCUPS from the combined GM12878 Hi-C data at 5kb resolution. For HiC-CUPS, from the 2, 993 loops where the two corresponding loci each contained a single CTCF-binding motif, we determined that 89.7% of motif pairs were convergent. There were 2,926 loops with a paired CTCF motif out of which 88.6% with convergent motifs for Mustache when the same number of loops were considered to that of HiCCUPS (i.e., top-*k*). When we used the fixed *q*-value threshold of 0.05 (resulting in a total of 18,068 loops), we obtained 5, 318 loops with a unique CTCF motifs on both ends, out of which 83.1% had convergent orientation. Mustache detected nearly two thousand additional convergent CTCF loops compared to HiCCUPS (a 45% increase) for GM12878, suggesting the existence of important structural loop that might have been missed by HiCCUPS.

Next, we compared the enrichment of structural markers CTCF, RAD21 and SMC3 [28] on the anchors of detected loops for both methods on the GM12878 cell line. First, we determined the set of loci (at 5kb resolution) that were involved in at least one reported chromatin loop for each method. Then for each interacting loci, we checked whether there was a midpoint of a ChIP-seq peak within 5kb and defined this locus as “overlapping” a CTCF or cohesin (SMC3 and RAD21) binding site if that was the case. This distance threshold for overlap detection was selected to replicate the findings of a similar analysis showing that 86-87% of interacting loci from HiCCUPS are “bound” by CTCF and cohesin (see Figure 6C [9]). When we compared the percentage of interacting loci overlapping ChIP-seq peaks according to the above criterion, these numbers were 82.5%, 82.7% and 80.7% for Mustache and 86.9%, 86.9% and 85.1% for HiCCUPS for CTCF, RAD21 and SMC3, respectively, when the number of loop calls from each method was fixed (top-*k* setting). These percentages were 74.5% for CTCF, 74.7% for RAD21 and 71.1% for SMC3 when we considered all Mustache loops at a *q*-value threshold of 0.05. These results suggest that interacting loci from Mustache have similar enrichment compared to those from HiCCUPS for overlapping binding of known looping related insulator proteins (top-*k* settings). When we consider interacting loci from over 10k additional loops reported by Mustache (q-value setting), we observe that nearly two thirds of them were also overlapping with CTCF and/or cohesin binding sites.

**Figure 3:**
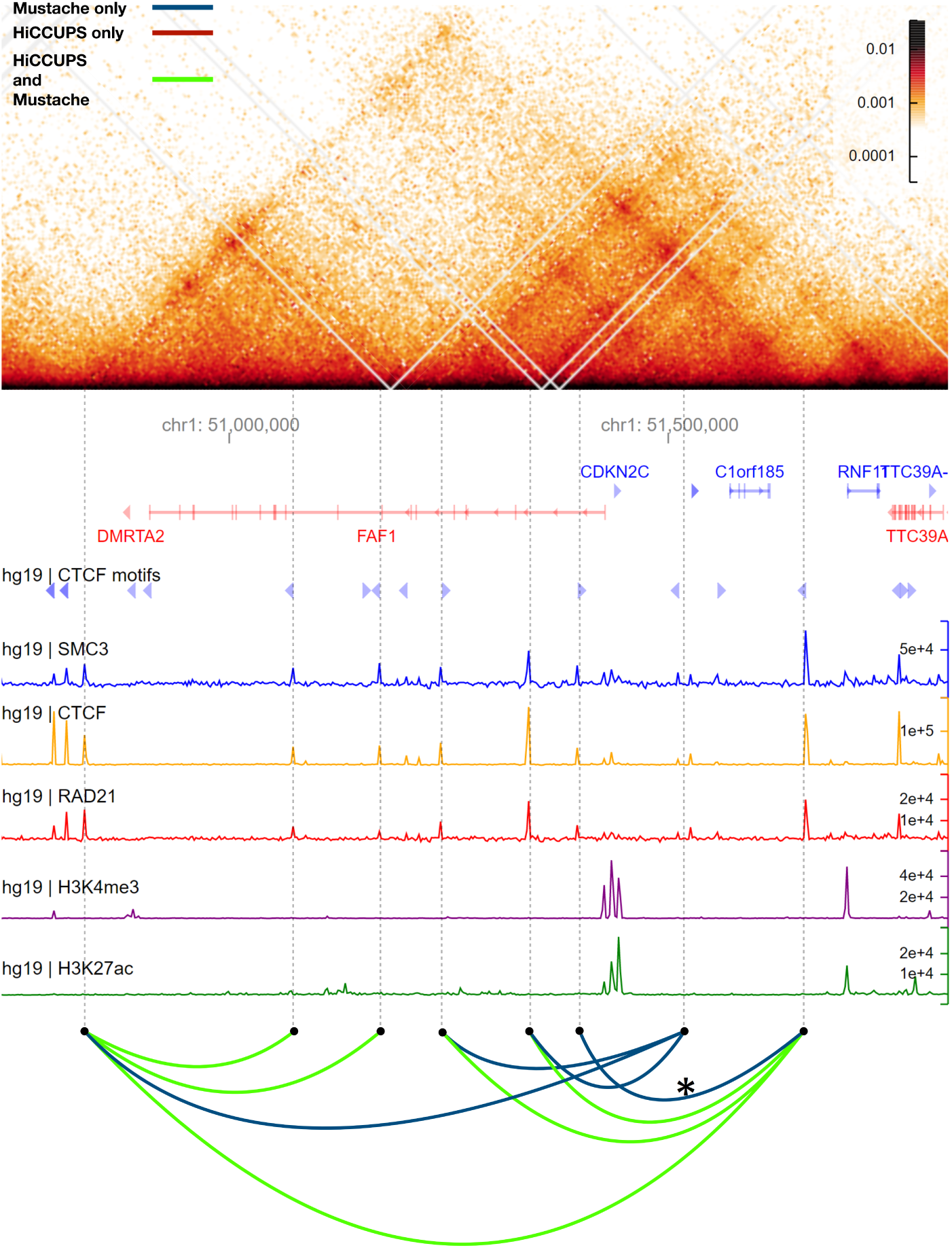
A comparison between Mustache and HiCCUPS reported loop calls in a region of chromosome 1 for the GM12878 cell line (50.75Mb−51.75Mb). The Hi-C contact map is rotated 45 degrees such that the main diagonal is horizontal (top). Below the contact map, we report genomic coordinates, gene annotations (genes on the negative strand are shown in red color), CTCF motifs and their orientation, ChIP-seq signals for SMC3, CTCF, RAD21, H3K4me3 and H3K27ac (coverage tracks plotted by HiGlass). The bottom row demonstrates loop calls as arcs connecting two loci with loops detected by both Mustache and HiCCUPS in green, only by Mustache in blue and only by HiCCUPS in red.

**Figure 4:**
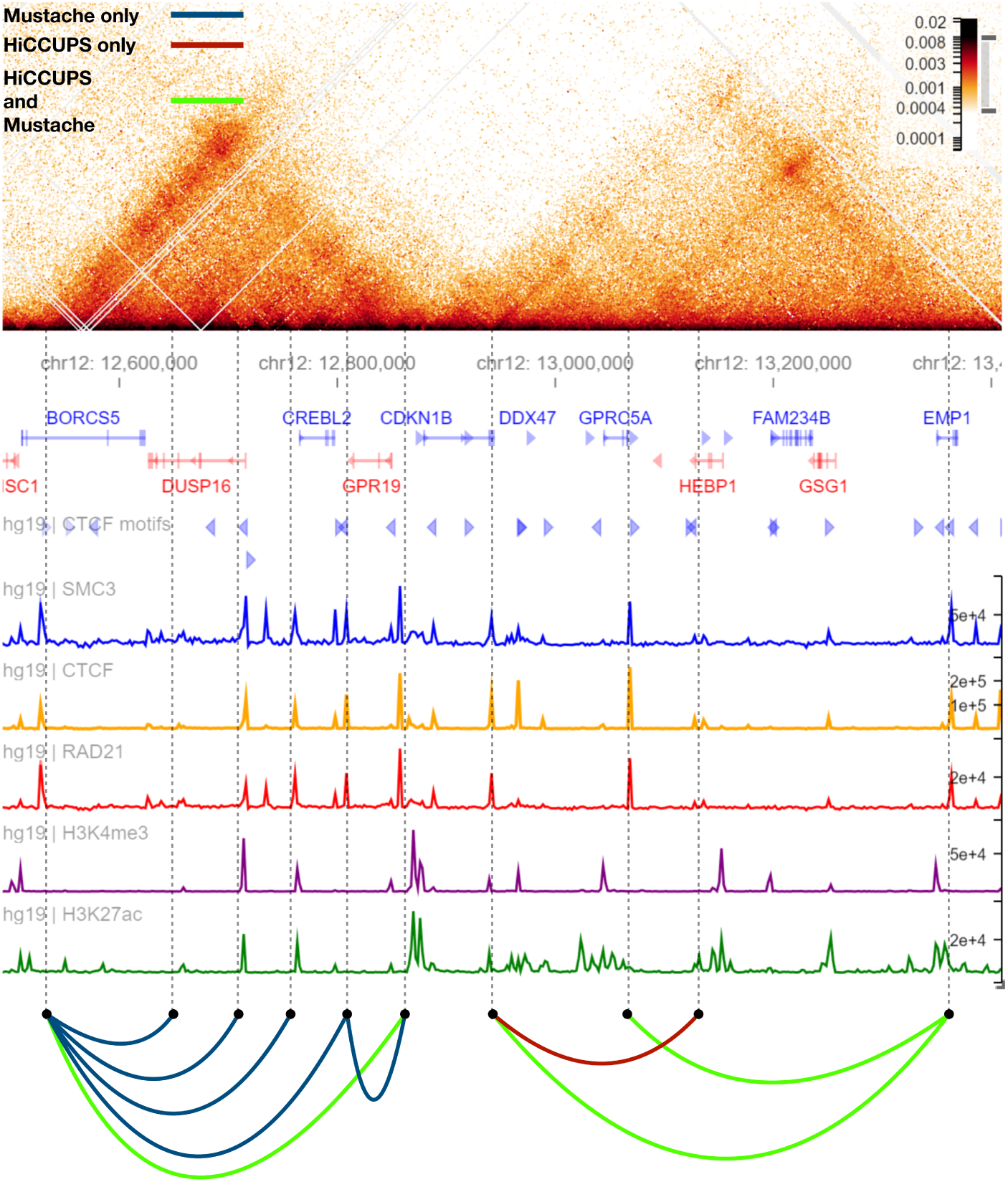
A comparison between Mustache and HiCCUPS reported loop calls in a region of chromosome 1 for the GM12878 cell line (12.5Mb−13.3Mb). The display order and color code are similar to that of Figure 3.

**Figure 5:**
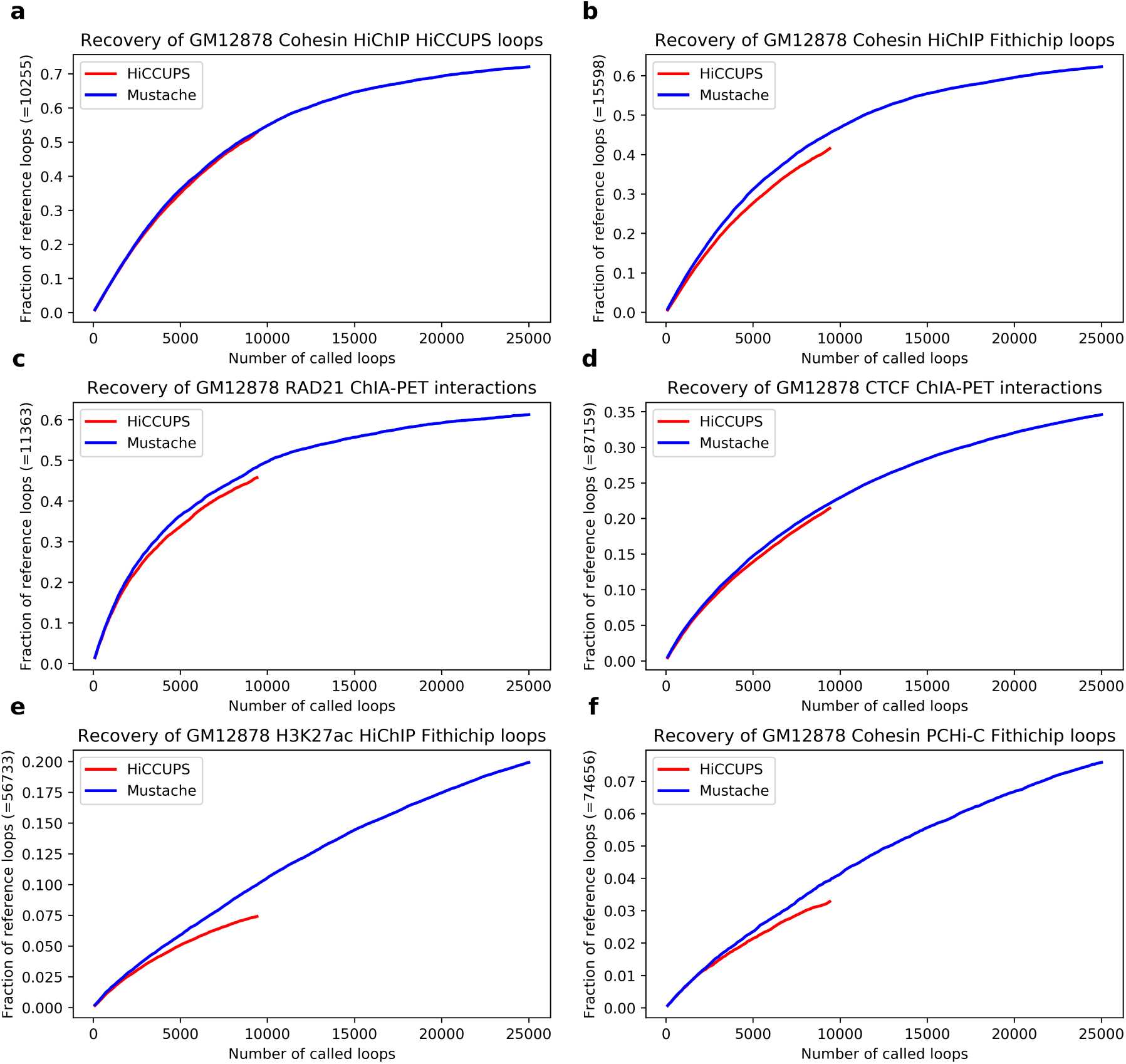
Comparison of the recovery of several difference reference loop sets by Mustache versus HiCCUPS applied on the 5kb resolution GM12878 Hi-C data. **(a)** GM12878 cohesin HiChIP HiCCUPS loops; **(b)** GM12878 cohesin HiChIP FitHiChIP loops; **(c)** GM12878 RAD21 ChIA-PET interactions; **(d)** GM12878 CTCF ChIA-PET interactions; **(e)** GM12878 H3K27ac HiChIP FitHiChIP loops; **(f)** GM12878 cohesin PCHiC FitHiChIP loops.

**Figure 6:**
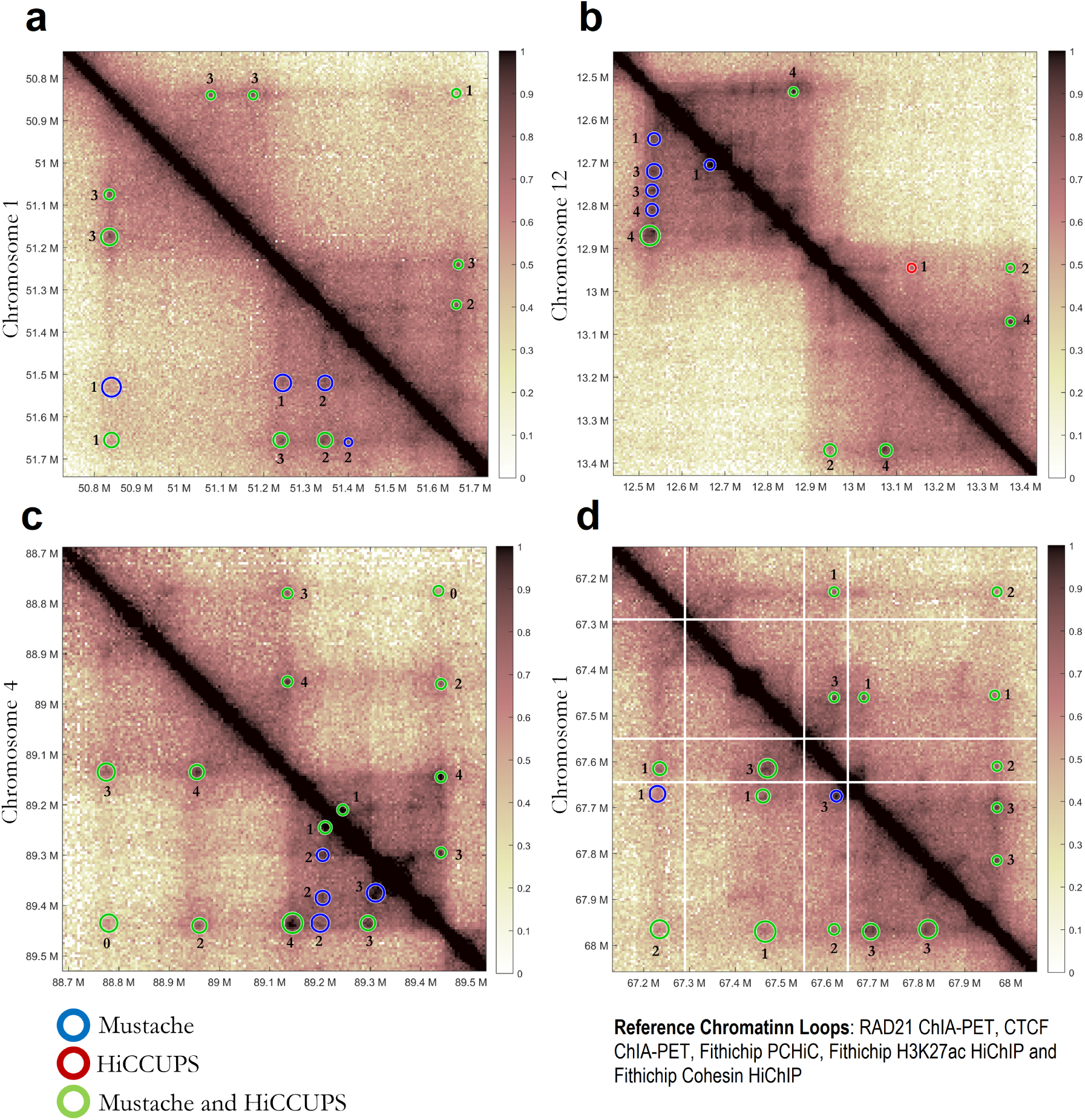
Four example regions showing Mustache and HiCCUPS reported loops. Shared loops between the two methods are denoted by green circles Method specific loops are shown using blue (Mustache-specific, lower triangular) and red circles (HiCCUPS-specific, upper triangular). The outputs of two methods using GM12878 Hi-C data are shown: **(a)** 50.75Mb-51.75Mb on chromosome 1, **(b)** 12.5Mb-13.4Mb on chromosome 12, **(c)** 88.7Mb-88.5Mb on chromosome 4, **(d)** 67.2Mb-68Mb on chromosome 1.

We, then, performed a similar overlap analysis, but this time by using promoter and enhancer annotations for the GM12878 cell lines as determined by ChromHMM [29]. More specifically, we counted the number of interacting pairs of loci, where one locus overlapped a promoter region, and the other locus overlapped an enhancer region. Here, we define the notion of overlap similar to ChIP-seq peaks discussed above, but instead using the midpoint of the ChromHMM annotation. The results showed that 38.9% (7019) of Mustache loops in GM12878 (q-value < 0.05) and 41.6% (3736) in K562 (q-value < 0.1) connected a promoter to an enhancer **(Supplementary Figure 4)**. The corresponding percentages for HiCCUPS were 37.8% (3576) of loops in GM12878 and 37% (2244) in K562. Similarly, 16.5% (2974) and 17.8% (1594) of Mustache loops, and 16.8% (1586) and 16.3% (988) percent of HiCCUPS loops connected a promoter to another promoter for GM12878 and K562 Hi-C data, respectively **(Supplementary Figure 4)**. These results suggested that the proportion of loops overlapping regulatory elements were similar for Mustache and HiCCUPS, with Mustache reporting a significantly higher number of such loops.

Lastly, we illustrate the relevance of Mustache’s improved detection power on some selected regions for the GM12878 Hi-C data, using HiGlass for visualization [30]. Figure 3 and Figure 4 provide a closer look at the regions 50.75Mb-51.75Mb on chromosome 1 and 12.5Mb-13.4Mb on chromosome 12, respectively. In these figures we included the 45-degree rotated Hi-C heatmaps, together with gene annotations, CTCF motifs and other relevant ChIP-seq signals. Figure 3 shows that Mustache detected all loops called by HiCCUPS for this locus (green arcs) and reported additional loops (blue arcs) that connected peaks of ChIP-seq signal from known correlates of looping such as CTCF and cohesin binding. For example, the loop denoted by “*” was detected only by Mustache, and it connected two loci that had strong peaks in all three structural signals and corresponded to a pair of loci with convergent CTCF motifs. The other three Mustache-only loops showed similar features of both cohesin and CTCF binding enrichment on each end. Figure 4 highlights another locus for which there were five Mustache-only loops, each with cohesin and CTCF binding peaks on each end, and three of them with a convergent CTCF pairing. In this genomic window, Mustache missed only one loop that was reported by HiCCUPS (red arc). Overall, these examples highlight that Mustache detects loops (missed by HiCCUPS) that have anchors enriched in genomic and epigenetic features known to be associated with chromatin loops.

### Mustache recovers a larger fraction of cell-type matched HiChIP and ChIA-PET loop calls

In the previous section we showed that Mustache detected additional chromatin loops (compared to HiCCUPS) that are supported by both genomic features (e.g., CTCF motifs) and functional (epi)genomics features (e.g., ChIP-seq, ChromHMM). Here, we compared Mustache and HiCCUPS loop calls using published ChIA-PET and HiChIP loops as reference. In this experiment, we computed the number of ChIA-PET and HiChIP loop calls recovered by Mustache and HiCCUPS from the 5kb resolution GM12878 data. Again, we used the matching criteria described in the reproducibility analysis to determine the overlap between two loop calls. Figure 5a shows the recovery plot on GM12878 cohesin HiChIP data [31]. The *x*-axis represents the number of Hi-C chromatin loops called by Mustache (blue) and HiCCUPS (red) sorted by their significance. We set HiCCUPS’ significance to be the median of the q-values over the four local filters. Mustache’s significance is the q-value reported, as described in Methods. The *y*-axis represents the percentage of the HiCCUPS loops from cohesin HiChIP data that were recovered by each method. Observe that HiCCUPS and Mustache on GM12878 Hi-C data had similar recovery when matching the number of loop calls. However, Mustache recovered more than 70% of all reference loops compared to less than 55% for HiCCUPS, even though the “reference” loops were previously determined using HiCCUPS on HiChIP data. We observed similar higher recovery for Mustache when we used FitHiChIP [21] to call cohesin HiChIP loops (Figure 5b). When we used ChIA-PET loops either from a RAD21 experiment (Figure 5c) [32] or from a CTCF experiment (Figure 5d) [33], Mustache provided a 10-15% improvement in recovery. For H3K27ac HiChIP data [34] and for promoter capture Hi-C (PCHiC) experiments [35] on the same cell line, the overall number of loop calls were several fold higher compared to either HiCCUPS or Mustache due to different types of looping that these experiments are designed to capture. However, Mustache still provided an improved recovery when either the same number of loops or the whole set of calls were considered (Figure 5e-f).

Taken all together, the genome-wide analyses described above showed that Mustache has better power in recovering loops identified from independent conformation capture experiments. To further elaborate on this point, we illustrate additional examples demonstrating that Mustache-specific loops are supported by other conformation capture data. Specifically, we asked for each Hi-C loop call from either method, how many of the five reference loop sets listed in Figure 6 have an overlapping loop call (number shown next to each circle), using the usual matching criteria defined above in the reproducibility analysis. The different radii of the circles illustrate the scale at which Mustache detected these loops.

Figure 6a and Figure 6b correspond to the two regions analyzed previously in Figure 3 and Figure 4, respectively. HiCCUPS and Mustache loops are represented by circles on the upper and lower diagonal matrix, respectively, with green color indicating common loops. For the 1.2Mb region on chromosome 1, Figure 6a highlights five shared loop calls (green) and four Mustache-specific loops as mentioned before. Each one of these four Mustache-specific loops were supported at least by one other reference set and two were supported by two independent lines of evidence. Figure 6b shows that Mustache detected five chromatin loops between an enhancer on the border of a topologically associating domains (TAD), four of which were specific to Mustache (upper-left corner of the shown contact map). Out of these four, three were supported by three or more reference sets. This locus also has one other Mustache-specific loop and one HiCCUPS-specific loop each of which was supported by one reference set. For the last two regions, one on chromosome 4 (Figure 6c) and one on chromosome 1 (Figure 6d), six Mustache-specific loops were supported by a several different reference sets (median number = 2). Observe that in Figure 6c and Figure 6d there were no HiCCUPS-specific loop. As expected, most of the detected loops fell inside or on the boundary of visible TADs for both methods. In general, there was high concordance between the two methods as indicated by 15 common loop calls in total. All these results taken together (along with the recovery analysis), suggest that the additional loops reported by Mustache are not false-positives, but they are very likely to be real looping events because they are supported by multiple lines of evidence with visible enrichment.

### Mustache detects loops that are consistent between Hi-C and Micro-C data

In this last set of experiments, we analyzed data from Micro-C conformation capture experiments [10]. Micro-C replaces the restriction enzyme used in Hi-C with micrococcal nuclease for digestion. Several groups recently used Micro-C to generate high-resolution contact maps of mouse and human cells highlighting its advantages over Hi-C and its potential to become an essential assay for nucleome studies [11, 12]. The objective here was to assess the performance of Mustache on Micro-C data and evaluate the consistency of Micro-C loops when compared to Hi-C loops on the same cell line. For this purpose, we used Mustache on Hi-C and Micro-C data (using the same parameters) for the HFFc6 human cell line [12]. Figure 7a shows that Mustache calls from Micro-C and Hi-C were largely consistent: 90% of Hi-C loops were also reported on the Micro-C data. However, Mustache identified over 20k more loops from Micro-C, which is more than double the number of loops detected from Hi-C contact maps. We then assessed how this overlap changed if only considered the top-*k* most significant Mustache loops from each dataset. Figure 7b shows the overlap fraction (y-axis) between the two datasets for different values of *k* ranging from 1k to 30k (x-axis). The overlap between Hi-C and Micro-C loops ranged between 60% to 74% with its maximum achieved when *k* is about 10k. This suggests that the ordering of loop candidates with respect to their significance is consistent between the two datasets, with Micro-C offering better statistical power for detection. In order to further understand this, we generated APA plots for loop calls from Hi-C and from Micro-C using the same contact maps for aggregation. Figure 7c-d show that the Micro-C loops had strikingly higher center pixel enrichment consistent with previous observations that Micro-C enriches signal-to-background ratio [12]. The reverse APA analysis (i.e., loop calls were taken from one dataset but the aggregation of contact patterns were computed using the other) showed that the Hi-C loops had a very striking center pixel enrichment in the Micro-C contact map, suggesting that Mustache accurately detected the correct looping locations also from Hi-C data but the local enrichment of the loops from Hi-C maps was not as strong as the loops obtained from Micro-C maps **(Supplementary Figure 5)**.

**Figure 7:**
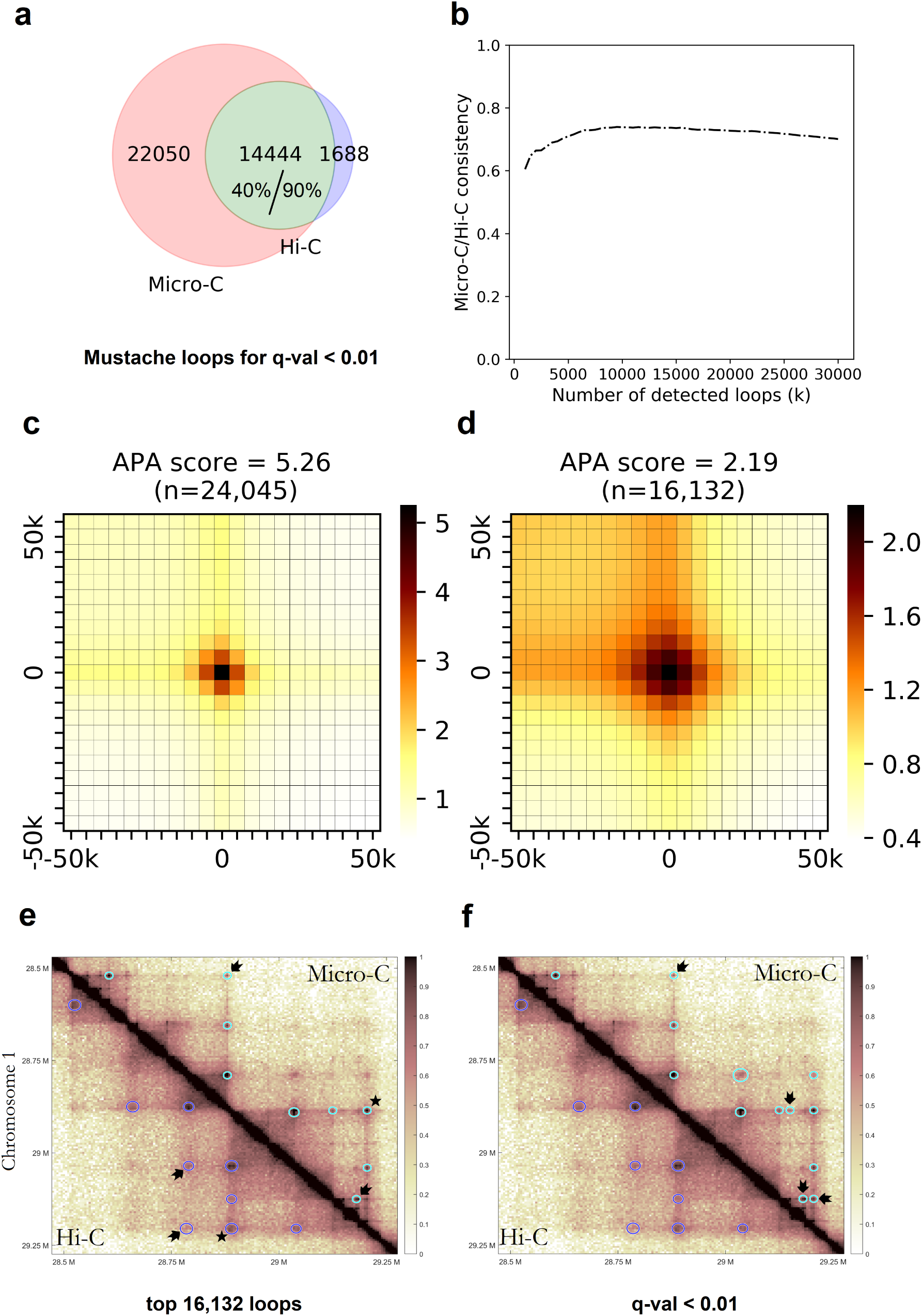
**(a)** The consistency between Mustache loops detected using Micro-C and Hi-C data in HFFc6 cell line using a fixed q-value threshold of 0.01, shown as a Venn diagram; **(b)** The consistency plot for Mustache results between Hi-C and Micro-C for the top-*k* reported interactions for each contact map. The APA plots for Mustache loops in HFFc6 cell line for **(c)** Micro-C and **(d)** Hi-C data. The APA score for the enrichment of center is reported above each plot. Mustache reported loops in Hi-C (lower triangular) and Micro-C (upper triangular) of HFFc6 for **(e)** the top 16,132 significant interactions, and **(f)** a fixed *q*-value threshold of 0.01. The loop call marked by a “*” was in common between Hi-C and Micro-C, but was detected at a smaller scale and has a stronger enrichment in Micro-C compared to Hi-C. The loops that are uniquely detected in either by Hi-C or by Micro-C are denoted by black arrows.

Finally, we visualized Hi-C and Micro-C loop calls from Mustache for an 800kb region on chromosome 1. Figure 7e shows the loops called by Mustache, when the tool was set to call the same number of loops on both datasets. Figure 7f shows loops called by Mustache using a significance threshold of 0.01. Hi-C loops are shown by blue circles on the lower triangular map, while Micro-C are shown by cyan circles in the upper triangular map. We used a common normalized color scale for the contact map visualization to account for sequencing depth differences. Observe in Figure 7e that the loop call marked by a “*” was in common between Hi-C and Micro-C, but was detected at a smaller scale and has a stronger enrichment in Micro-C compared to Hi-C. This example is consistent with the genome-wide patterns we summarized above using APA analysis. Also observe in Figure 7e that despite the fact that the same number of loops were called for both data types, Mustache had different priorities in selecting loops due to the underlying differences between Hi-C and Micro-C outputs. This is also visible from the contact maps: loops that were uniquely detected in Hi-C or Micro-C are denoted by black arrows. When a fixed *q*-value threshold is used, Figure 7f shows that Mustache called more loops from Micro-C compared with Hi-C including the two “Hi-C-specific” loops when top-*k* loops were considered (Figure 7e). These additional Micro-C loops had strong local enrichment only visible on the Micro-C contact maps. This is possibly due to dilution of contact enrichment for some loops into multiple pixels for Hi-C in contrast to their concentration on a specific pixel when a higher resolution digestion system is used such as Micro-C. Taken together, all these experimental results show Mustache provides highly consistent results on Micro-C and Hi-C data, but can leverage the unique strengths of each protocol to discover loops that are specifically enriched by that essay.

## Conclusion

The rapid adoption of Hi-C and its variants has fueled efforts in the development of computational tools to study 3D chromatin organization including chromosome compartments, topologically associating domains and chromatin loops [8, 36, 9, 37, 38]. Here we presented a novel method for the identification of chromatin loops that uses a scale-space representation of Hi-C and Micro-C contact maps. Experimental results show that loops detected by our method are not only enriched with respect to their global expectation (as quantified by methods developed by us and others [17, 18, 20]) but they also strongly correlated to known signals such as convergent CTCF binding sites or the boundaries of topologically associating domains [9]. Our novel multi-scale approach allows Mustache to account for the dependencies among neighboring pixels (both spatially and across resolutions) of the contact map, which are ignored by methods that use global background estimates for significance calculation.

Our experimental results on Hi-C and Micro-C contact maps show that Mustache robustly identifies looping events including those reported by HiCCUPS [9], as well as those detected by independent chromatin conformation capture experiments, including ChIA-PET and HiChIP. Mustache has several advantages over HiCCUPS, namely **i** it is very time efficient, and its implementation does not require GPUs (e.g., it takes only a few minutes per chromosome for 5kb resolution human Hi-C maps on a standard laptop with CPU), **ii** it has better reproducibility of loop calls from replicate experiments, **iii** it provides higher statistical power that results in a higher number of chromatin loops (additional loops are strongly supported by genomic and epigenetic features), and **iv** it produces highly consistent loop calls when comparing Hi-C and Micro-C data, allowing a robust comparative analysis of the two methods. In summary, Mustache represents a significant improvement over the state-of-the-art for the analysis of chromatin organization from high-resolution Hi-C and Micro-C contact maps.

## Availability of data and materials

A Python implementation of Mustache is freely available at https://github.com/ay-lab/mustache. This implementation works with commonly used file formats for Hi-C data including “.hic” and “.cool” formats. The accession numbers of all data used in this manuscript are available in supplementary file, Table S1. All reported chromatin loops as well as the ChIP-seq data used in the manuscript are available on Washu Epigenome browser [39]. To load the data in the Washu Epigenome browser (http://epigenomegateway.wustl.edu) please provide the following URL after selecting “Explore the new browser - human - hg19” then going to the “Tracks - Custom Tracks - Add Custom Data Hub”. https://informaticsdata.liai.org/BioAdHoc/Groups/vd-ay/abbas/epgBrowser/Mustache_all_WashU.json

## Author contributions

A. R. A., S. L. and F.A. conceptualized the method. A. R. A. implemented the method and applied it to public data. H. T. G. wrote the initial Python code and A. R. A. revised and finalized this implementation. A. R. A. and F. A. interpreted the experimental results. A. R. A., F. A. and S. L. wrote the manuscript with input from H. T. G. All authors read and approved the final version of the manuscript.

## Acknowledgements

We would like to acknowledge all members of Lonardi Lab at UC Riverside and Ay Lab at La Jolla Institute for Immunology for their valuable feedback. This work was funded in part by NIH grant R35-GM128938 to F.A. and NSF grant IIS-1814359 to S.L.

## References

[1] Pederson, T.: Chromatin structure and the cell cycle. Proc. Natl. Acad. Sci. U. S. A. 69(8), 2224–2228 (1972)

[2] Dixon, J.R., Xu, J., Dileep, V., Zhan, Y., Song, F., Le, V.T., Yardımcı, G.G., Chakraborty, A., Bann, D.V., Wang, Y., et al.: Integrative detection and analysis of structural variation in cancer genomes. Nat Genet. 50(10), 1388–1398 (2018)

[3] Dileep, V., Ay, F., Sima, J., Vera, D.L., Noble, W.S., Gilbert, D.M.: Topologically associating domains and their long-range contacts are established during early G1 coincident with the establishment of the replication-timing program. Genome Res. 25(8), 1104–1113 (2015)

[4] Beagrie, R.A., Pombo, A.: Cell cycle: Continuous chromatin changes. Nature 547(7661), 34–35 (2017)

[5] Bonev, B., Cavalli, G.: Organization and function of the 3D genome. Nat Rev. Genet. 17(11), 661–678 (2016)

[6] Zheng, H., Xie, W.: The role of 3D genome organization in development and cell differentiation. Nat Rev Mol Cell Biol. 20(9), 535–550 (2019)

[7] Marchal, C., Sima, J., Gilbert, D.M.: Control of DNA replication timing in the 3D genome. Nat Rev Mol Cell Biol. 20, 1–17 (2019)

[8] Lieberman-Aiden, E., van Berkum, N.L., Williams, L., Imakaev, M., Ragoczy, T., Telling, A., Amit, I., Lajoie, B.R., Sabo, P.J., Dorschner, M.O., Sandstrom, R., Bernstein, B., Bender, M.A., Groudine, M., Gnirke, A., Stamatoyannopoulos, J., Mirny, L.A., Lander, E.S., Dekker, J.: Comprehensive mapping of long-range interactions reveals folding principles of the human genome. Science 326(5950), 289–293 (2009)

[9] Rao, S.S.P., Huntley, M.H., Durand, N.C., Stamenova, E.K., Bochkov, I.D., Robinson, J.T., Sanborn, A.L., Machol, I., Omer, A.D., Lander, E.S., Aiden, E.L.: A 3D map of the human genome at kilobase resolution reveals principles of chromatin looping. Cell 159(7), 1665–1680 (2014)

[10] Hsieh, T.-H.S., Weiner, A., Lajoie, B., Dekker, J., Friedman, N., Rando, O.J.: Mapping nucleosome resolution chromosome folding in yeast by micro-c. Cell 162(1), 108–119 (2015)

[11] Hansen, A.S., Hsieh, T.-H.S., Cattoglio, C., Pustova, I., Saldaña-Meyer, R., Reinberg, D., Darzacq, X., Tjian, R.: Distinct classes of chromatin loops revealed by deletion of an RNA-binding region in CTCF. Mol Cell. 76(3), 395–411 (2019)

[12] Krietenstein, N., Abraham, S., Venev, S.V., Abdennur, N., Gibcus, J., Hsieh, T.-H.S., Parsi, K.M., Yang, L., Maehr, R., Mirny, L.A., Dekker, J., Rando, O.J.: Ultrastructural details of mammalian chromosome architecture. bioRxiv (2019). doi: 10.1101/639922

[13] Bonev, B., Mendelson Cohen, N., Szabo, Q., Fritsch, L., Papadopoulos, G.L., Lubling, Y., Xu, X., Lv, X., Hugnot, J.-P., Tanay, A., Cavalli, G.: Multiscale 3D genome rewiring during mouse neural development. Cell 171(3), 557–57224 (2017)

[14] Fudenberg, G., Abdennur, N., Imakaev, M., Goloborodko, A., Mirny, L.A.: Emerging evidence of chromosome folding by loop extrusion. In: Cold Spring Harbor Symposia on Quantitative Biology, vol. 82, pp. 45–55 (2017). Cold Spring Harbor Laboratory Press

[15] Davidson, I.F., Bauer, B., Goetz, D., Tang, W., Wutz, G., Peters, J.-M.: DNA loop extrusion by human cohesin. Science 366(6471), 1338–1345 (2019)

[16] Sanborn, A.L., Rao, S.S., Huang, S.-C., Durand, N.C., Huntley, M.H., Jewett, A.I., Bochkov, I.D., Chinnappan, D., Cutkosky, A., Li, J., et al.: Chromatin extrusion explains key features of loop and domain formation in wild-type and engineered genomes. Proc. Natl. Acad. Sci. U. S. A.” 112(47), 6456–6465 (2015)

[17] Ay, F., Bailey, T.L., Noble, W.S.: Statistical confidence estimation for Hi-C data reveals regulatory chromatin contacts. Genome Res. 24(6), 999–1011 (2014)

[18] Carty, M., Zamparo, L., Sahin, M., González, A., Pelossof, R., Elemento, O., Leslie, C.S.: An integrated model for detecting significant chromatin interactions from high-resolution hi-c data. Nat Commun. 8, 15454 (2017)

[19] Zufferey, M., Tavernari, D., Oricchio, E., Ciriello, G.: Comparison of computational methods for the identification of topologically associating domains. Genome Biol. 19(1), 217 (2018)

[20] Forcato, M., Nicoletti, C., Pal, K., Livi, C.M., Ferrari, F., Bicciato, S.: Comparison of computational methods for Hi-C data analysis. Nat Methods 14(7), 679 (2017)

[21] Bhattacharyya, S., Chandra, V., Vijayanand, P., Ay, F.: Identification of significant chromatin contacts from HiChIP data by FitHiChIP. Nat Commun. 10(1), 4221 (2019). doi: 10.1038/s41467-019-11950-y

[22] Kaul, A., Bhattacharyya, S., Ay, F.: Identifying statistically significant chromatin contacts from Hi-C data with FitHiC2. Nat Prot. (2020). doi: 10.1038/s41596-019-0273-0

[23] Dekker, J., Belmont, A.S., Guttman, M., Leshyk, V.O., Lis, J.T., Lomvardas, S., Mirny, L.A., O’Shea, C.C., Park, P.J., Ren, B., Politz, J.C.R., Shendure, J., Zhong, S., 4D Nucleome Network: The 4D nucleome project. Nature 549(7671), 219–226 (2017)

[24] Lindeberg, T.: Scale-Space Theory in Computer Vision. Springer, Boston, MA (1994)

[25] Lowe, D.G.: Distinctive image features from Scale-Invariant keypoints. Int. J. Comput. Vis. 60(2), 91–110 (2004)

[26] Lowe, D.G.: Object recognition from local scale-invariant features. In: Proceedings of the Seventh IEEE International Conference on Computer Vision, vol. 2, pp. 1150–11572 (1999)

[27] Benjamini, Y., Hochberg, Y.: Controlling the false discovery rate: A practical and powerful approach to multiple testing. J. R. Stat. Soc. Series B Stat. Methodol. 57(1), 289–300 (1995)

[28] Phanstiel, D.H., Boyle, A.P., Heidari, N., Snyder, M.P.: Mango: a bias-correcting ChIA-PET analysis pipeline. Bioinformatics 31(19), 3092–3098 (2015)

[29] Ernst, J., Kellis, M.: ChromHMM: automating chromatin-state discovery and characterization. Nat Methods 9(3), 215–216 (2012)

[30] Kerpedjiev, P., Abdennur, N., Lekschas, F., McCallum, C., Dinkla, K., Strobelt, H., Luber, J.M., Ouellette, S.B., Azhir, A., Kumar, N., et al.: HiGlass: web-based visual exploration and analysis of genome interaction maps. Genome Biol. 19(1), 125 (2018)

[31] Mumbach, M.R., Rubin, A.J., Flynn, R.A., Dai, C., Khavari, P.A., Greenleaf, W.J., Chang, H.Y.: HiChIP: efficient and sensitive analysis of protein-directed genome architecture. Nat Methods 13(11), 919 (2016)

[32] Heidari, N., Phanstiel, D.H., He, C., Grubert, F., Jahanbani, F., Kasowski, M., Zhang, M.Q., Snyder, M.P.: Genome-wide map of regulatory interactions in the human genome. Genome Res. 24(12), 1905–1917 (2014)

[33] Tang, Z., Luo, O.J., Li, X., Zheng, M., Zhu, J.J., Szalaj, P., Trzaskoma, P., Magalska, A., Wlodarczyk, J., Ruszczycki, B., Michalski, P., Piecuch, E., Wang, P., Wang, D., Tian, S.Z., Penrad-Mobayed, M., Sachs, L.M., Ruan, X., Wei, C.-L., Liu, E.T., Wilczynski, G.M., Plewczynski, D., Li, G., Ruan, Y.: CTCF-Mediated human 3D genome architecture reveals chromatin topology for transcription. Cell 163(7), 1611–1627 (2015)

[34] Mumbach, M.R., Satpathy, A.T., Boyle, E.A., Dai, C., Gowen, B.G., Cho, S.W., Nguyen, M.L., Rubin, A.J., Granja, J.M., Kazane, K.R., Wei, Y., Nguyen, T., Greenside, P.G., Corces, M.R., Tycko, J., Simeonov, D.R., Suliman, N., Li, R., Xu, J., Flynn, R.A., Kundaje, A., Khavari, P.A., Marson, A., Corn, J.E., Quertermous, T., Greenleaf, W.J., Chang, H.Y.: Enhancer connectome in primary human cells identifies target genes of disease-associated DNA elements. Nat Genet. 49(11), 1602–1612 (2017)

[35] Mifsud, B., Tavares-Cadete, F., Young, A.N., Sugar, R., Schoenfelder, S., Ferreira, L., Wingett, S.W., Andrews, S., Grey, W., Ewels, P.A., et al.: Mapping long-range promoter contacts in human cells with high-resolution capture Hi-C. Nat Genet. 47(6), 598 (2015)

[36] Dixon, J.R., Selvaraj, S., Yue, F., Kim, A., Li, Y., Shen, Y., Hu, M., Liu, J.S., Ren, B.: Topological domains in mammalian genomes identified by analysis of chromatin interactions. Nature 485(7398), 376–380 (2012)

[37] Ay, F., Noble, W.S.: Analysis methods for studying the 3D architecture of the genome. Genome Biol. 16(1), 183 (2015)

[38] Chakraborty, A., Ay, F.: The role of 3D genome organization in disease: From compartments to single nucleotides. Semin Cell Dev Biol 90, 104–113 (2019)

[39] Li, D., Hsu, S., Purushotham, D., Sears, R.L., Wang, T.: WashU Epigenome Browser update 2019. Nucleic Acids Res. 47(W1), 158–165 (2019)

