## Supplementary Information for "Mustache: Multi-scale Detection of Chromatin Loops from Hi-C and Micro-C Maps using Scale-Space Representation"

February 24, 2020

| **Accession** | **Data Type** | **Cell Type** | **Factor** | **Reference** |
|---|---|---|---|---|
| GSM733752 | ChIP-seq | GM12878 | CTCF | ENCODE[1] |
| GSM935376 | ChIP-seq | GM12878 | Cohesin (SMC3) | ENCODE[1] |
| GSM803416 | ChIP-seq | GM12878 | RAD21 | ENCODE[1] |
| GSE80820 | HiChIP | GM12878 | Cohesin (SMC1A) | Mumbach et al. 2016 [5] |
| GSE101498 | HiChIP | GM12878 | H3K27ac | Mumbach et al. 2017 [6] |
| GSM1872886 | ChIA-PET | GM12878 | CTCF | Tang et al. 2015 [8] |
| GSM1436265 | ChIA-PET | GM12878 | Cohesin (RAD21) | Heidari et al. 2014 [2] |
| GSE63525 | Hi-C | GM12878 | None | Rao et al. 2014 [7] |
| GSE63525 | Hi-C | K562 | None | Rao et al. 2014 [7] |
| E-MTAB-2323 | PCHiC | GM12878 | None | Mifsud et al. [4] |
| 4DNES2R6PUEK | Hi-C | HFFc6 | None | Krietenstein et al. (2019)[3] |
| 4DNESYTWHUH6 | Micro-c | HFFc6 | None | Krietenstein et al. (2019)[3] |

Table 1: **List of datasets used in the current study.**

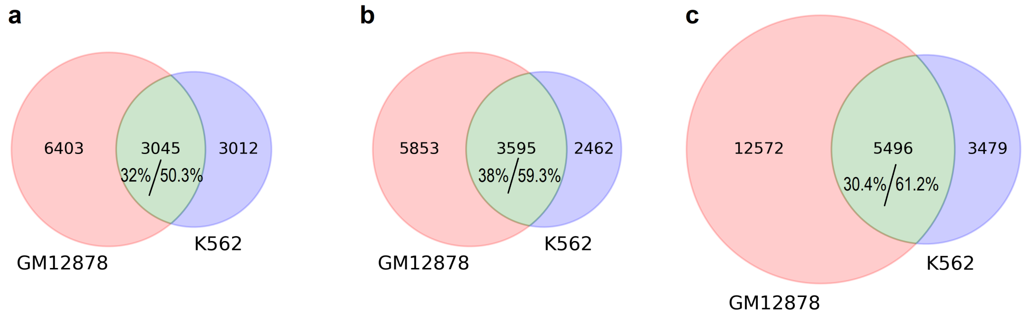


Figure 1: Reported chromatin loops in GM12878 and K562 cell lines and the overlap between them. The overlap is shown in green and the percentages of overlap with respect to each set are reported separately; **(a)** HiCCUPS reported loops in GM12878 and K562; **(b)** Mustache reported loops in GM12878 and K562 (with the same number of loops as in **(a)**); **(c)** Mustache reported loops in GM12878 and K562 for $q$-value threshold of 0.05 for GM12878 and $q$-value threshold of 0.1 for K562.

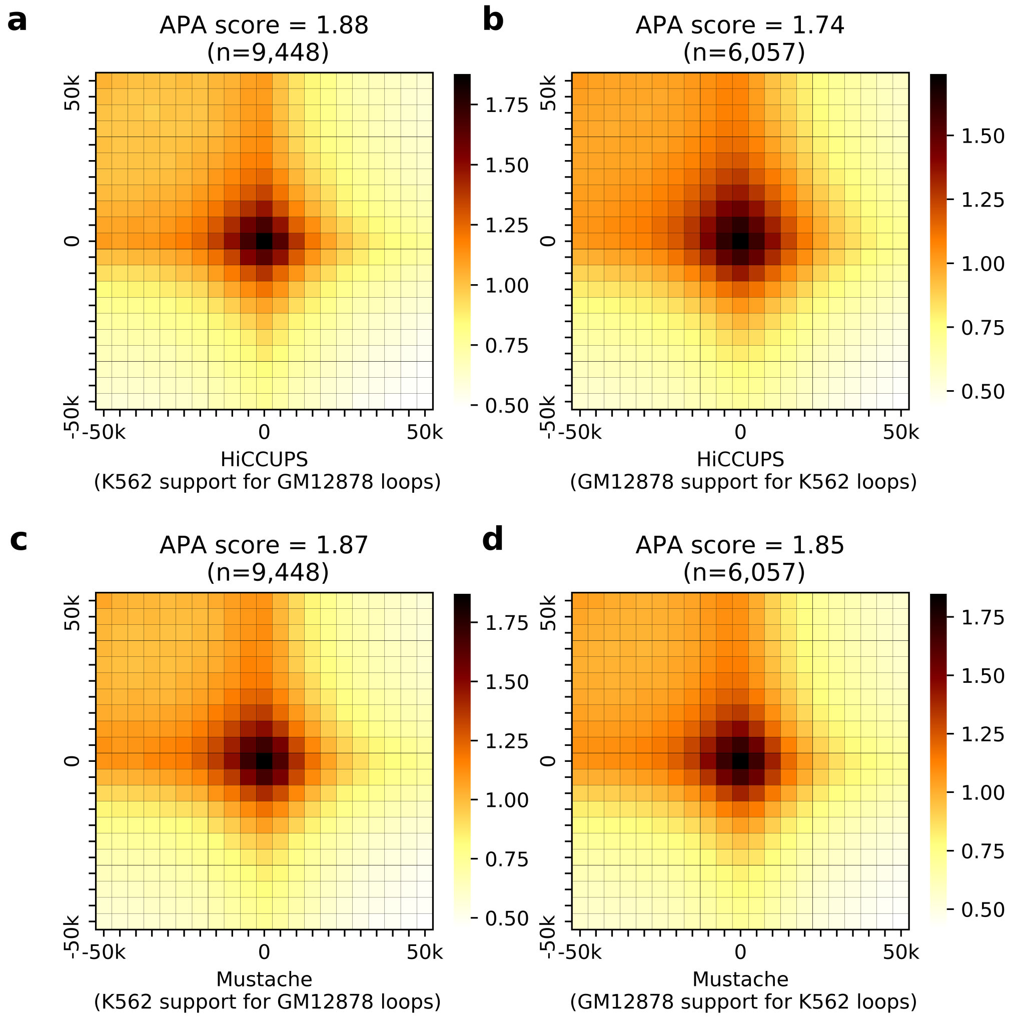


Figure 2: APA plots for HiCCUPS Hi-C loops in **(c)** GM12878 cell line supported by K562 contact maps, and **(d)** in K562 cell line supported by GM12878 contact maps; APA plots for Mustache Hi-C loops in **(c)** GM12878 cell line supported by K562 contact maps, and **(d)** in K562 cell line supported by GM12878 contact maps.

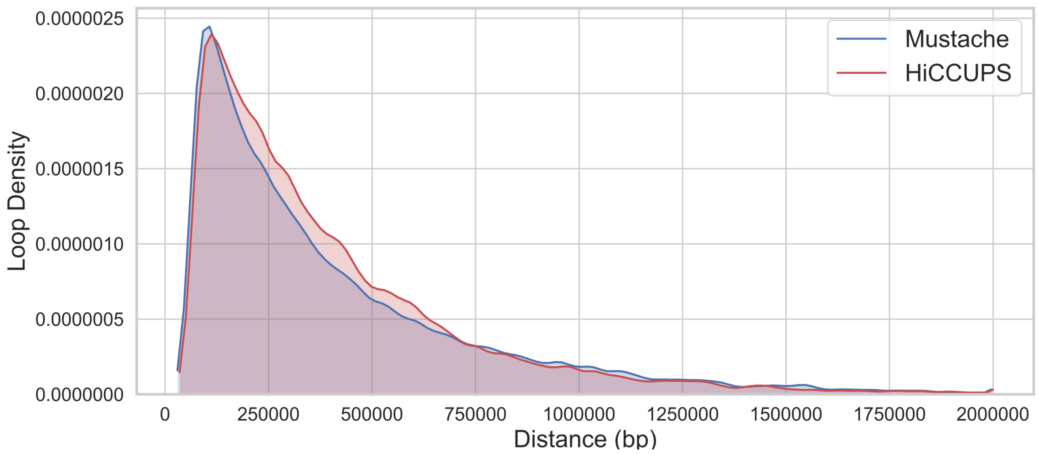


Figure 3: Genomic distance distribution between the loci of chromatin loops detected by MUSTACHE and HICCUPS in GM12878 cell line.

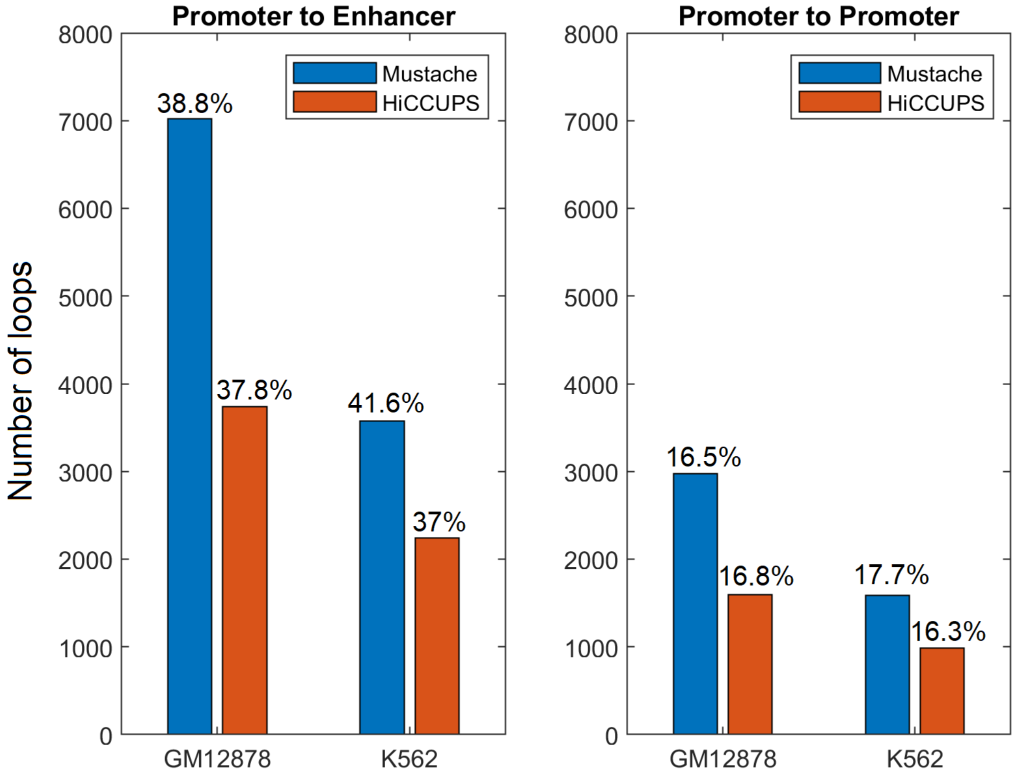


Figure 4: The number of chromatin interactions detected by MUSTACHE and HiCCUPS that connect promoters to enhancers and promoters to promoters (according to ChromHMM chromatin states). The percentage of all loops called by each method, is reported above each bar; **(a)** The number of chromatin loops detected by MUSTACHE and HiCCUPS that connect promoters to enhancers in cell lines GM12878 and K562. **(b)** The number of chromatin loops detected by MUSTACHE and HiCCUPS that connect promoters to promoters in cell lines GM12878 and K562.

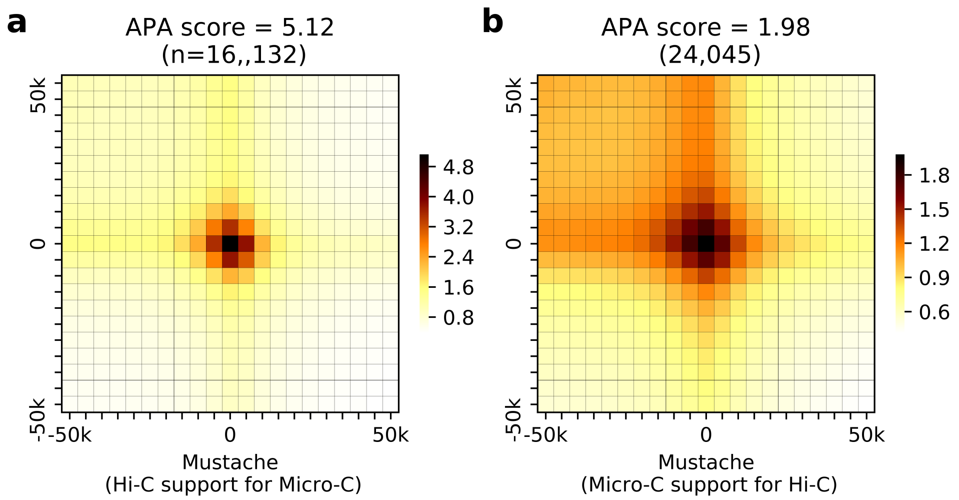


Figure 5: APA plots for Mustache reported loops in HFFc6 cell line **(a)** detected using Hi-C data, supported by Micro-C contact maps, and **(b)** detected using Micro-C data, supported by Hi-C contact maps.

### References


[1] ENCODE Project Consortium et al. An integrated encyclopedia of dna elements in the human genome. *Nature*, 489(7414):57, 2012.

[2] Nastaran Heidari, Douglas H Phanstiel, Chao He, Fabian Grubert, Fereshteh Jahanbani, Maya Kasowski, Michael Q Zhang, and Michael P Snyder. Genome-wide map of regulatory interactions in the human genome. *Genome Res.*, 24(12):1905–1917, December 2014.

[3] Nils Krietenstein, Sameer Abraham, Sergey V. Venev, Nezar Abdennur, Johan Gibcus, Tsung-Han S. Hsieh, Krishna Mohan Parsi, Liyan Yang, René Maehr, Leonid A. Mirny, Job Dekker, and Oliver J. Rando. Ultrastructural details of mammalian chromosome architecture. *bioRxiv*, 2019.

[4] Borbala Mifsud, Filipe Tavares-Cadete, Alice N Young, Robert Sugar, Stefan Schoenfelder, Lauren Ferreira, Steven W Wingett, Simon Andrews, William Grey, Philip A Ewels, et al. Mapping long-range promoter contacts in human cells with high-resolution capture hi-c. *Nature genetics*, 47(6):598, 2015.

[5] Maxwell R Mumbach, Adam J Rubin, Ryan A Flynn, Chao Dai, Paul A Khavari, William J Greenleaf, and Howard Y Chang. Hichip: efficient and sensitive analysis of protein-directed genome architecture. *Nature methods*, 13(11):919, 2016.

[6] Maxwell R Mumbach, Ansuman T Satpathy, Evan A Boyle, Chao Dai, Benjamin G Gowen, Seung Woo Cho, Michelle L Nguyen, Adam J Rubin, Jeffrey M Granja, Katelynn R Kazane, Yuning Wei, Trieu Nguyen, Peyton G Greenside, M Ryan Corces, Josh Tycko, Dimitre R Simeonov, Nabeela Suliman, Rui Li, Jin Xu, Ryan A Flynn, Anshul Kundaje, Paul A Khavari, Alexander Marson, Jacob E Corn, Thomas Quertermous, William J Greenleaf, and Howard Y Chang. Enhancer connectome in primary human cells identifies target genes of disease-associated DNA elements. *Nat. Genet.*, 49(11):1602–1612, November 2017.

[7] Suhas S P Rao, Miriam H Huntley, Neva C Durand, Elena K Stamenova, Ivan D Bochkov, James T Robinson, Adrian L Sanborn, Ido Machol, Arina D Omer, Eric S Lander, and Erez Lieberman Aiden. A 3D map of the human genome at kilobase resolution reveals principles of chromatin looping. *Cell*, 159(7):1665–1680, 18 December 2014.

[8] Zhonghui Tang, Oscar Junhong Luo, Xingwang Li, Meizhen Zheng, Jacqueline Jufen Zhu, Przemyslaw Szalaj, Pawel Trzaskoma, Adriana Magalska, Jakub Wlodarczyk, Blazej Ruszczycki, Paul Michalski, Emaly Piecuch, Ping Wang, Danjuan Wang, Simon Zhongyuan Tian, May Penrad-Mobayed, Laurent M Sachs, Xiaoan Ruan, Chia-Lin Wei, Edison T Liu, Grzegorz M Wilczynski, Dariusz Plewczynski, Guoliang Li, and Yijun Ruan. CTCF-Mediated human 3D genome architecture reveals chromatin topology for transcription. *Cell*, 163(7):1611–1627, December 2015.
